# In-vivo protein nitration and de-nitration facilitate *Vibrio cholerae* cell survival under anaerobic condition: Consequences of Nitrite induced protein nitration

**DOI:** 10.1101/2021.03.19.436110

**Authors:** Sourav Kumar Patra, Nilanjan Sinha, Subhamoy Chakraborty, Ayantika Sengupta, Souvik Roy, Sanjay Ghosh

## Abstract

Protein tyrosine nitration (PTN), a highly selective post translational modification, occurs in both prokaryotic and eukaryotic cells under nitrosative stress^1^. It is reported that the activities of many proteins are altered due to PTN^2^. PTN is found to be associated with many pathophysiological conditions like neurodegenerative and cardiac diseases etc.^3^. However, its physiological function is not yet clear. Like all other gut pathogens *Vibrio cholerae* also faces nitrosative stress in the gut environment which makes its proteome more vulnerable to PTN. Here, we report for the first time in-vivo PTN in *V. cholerae*. We show that in-vivo protein nitration is nitrite dependent and nitration-denitration phenomenon actually facilitates *V. cholerae* cell survival in anaerobic or hypoxic condition. In our study, we found that the extent of in-vivo nitration is negatively correlated with the intracellular nitrite content and maximum nitration occurs during log phase of *V. cholerae*. Most interestingly, a significant denitration was associated with increase in intracellular nitrate content during anaerobic incubation of aerobically grown late log phase cultures. In-vivo nitration could provide an avenue for toxic nitrite storage and nitrosative stress tolerance mechanism in many gut pathogens, whereas denitration could supply nitrate for cell survival in anaerobic nitrate deficient environment.

---

Protein Tyrosine Nitration (PTN) is an in-vivo post translational modification which has been reported in both prokaryotic and eukaryotic cells^1^. PTN is found to occur in many pathophysiological settings like cardiovascular diseases, lung diseases, neurodegenerative diseases and diabetes under inflammatory conditions where cells face nitrosative stress caused by excess nitric oxide and peroxynitrite^4,5,6^. In PTN, Tyrosine is modified in the 3-position of the phenolic ring through the addition of a nitro group. It is believed that tyrosine nitration involves a two-step process where the initial step is the oxidation of the phenolic ring of Tyr to yield the one electron oxidation product, Tyr radical (Tyr·). The second step involves the addition of NO_2_ to the Tyr · in a radical termination reaction^7^. There are two proximal nitrating agents that account for nitration in vivo. One nitrating agent is peroxynitrite which is formed by the fast reaction between nitric oxide (NO) and superoxide (O_2_·). The other proximal nitrating agents involve heme peroxidases such as myeloperoxidase or eosinophil peroxidase in the presence of hydrogen peroxide (H_2_O_2_) and nitrite (NO_2_)^8^. Based on the in vivo and in vitro data it is observed that PTN is a highly selective process in which the local structural environment of specific tyrosine residues governs the selectivity^9^. Protein tyrosine nitration caused several types of responses in terms of activity. In most cases PTN inhibits enzyme activity, in some cases activity remained unchanged and few reports show activation of the proteins or enzymes^2, 10, 11, 12^. However, the physiological role of PTN is yet to be identified.

Clinical studies have showed that serum and urine of the patients suffering from cholera have an increased NO metabolite level and increased inducible nitric oxide synthases (iNOS) expression in their small intestine during infection of *V. cholerae*^13^. Thus, *V. cholerae* encounters NO during infection in human. During our investigation we surprisingly observed in-vivo protein tyrosine nitration under different growth phases of *V. cholerae* grown separately in rich media as well as in minimal media using monoclonal nitrotyrosine antibody and LC-ESI-MS/MS analysis. A series of proteins were found to be nitrated at single tyrosine or multiple tyrosine residues. In the present study, we discovered a role of nitrite in in-vivo protein nitration of *V. cholerae*. During our study we surprisingly found de-nitration phenomenon of *V. cholerae* proteome when aerobically grown late log phase cells were incubated under anaerobic condition. We checked the cellular redox status of anaerobically incubated cells along with NO_3_ and NO_2_ content of the cytosol. Where, a significant increase in intracellular nitrate content was observed along with a controlled oxidative environment i.e. low GSH/GSSG. Based upon our data, it is conceivable that in-vivo nitration could provide a mechanism to store toxic nitrite inside cellular proteome without affecting *V. cholerae*’s growth. At the same time we found that nitration and de-nitration phenomenon is a cyclical process, which is essential for cellular respiration and cell survival in nutrient deprived anaerobic or hypoxic condition. In-vivo nitration could also be an impressive evolutionary adapted mechanism for nitrosative stress tolerance as it minimizes the probability of further nitration of proteome.

## Results

### Growth phase specific in vivo protein nitration profile of *V. cholerae* cells grown in rich media under aerobic condition

Being a gut pathogen *V. cholerae* continuously face nitrosative stress in the local environment which is supported by the production of excess NO and its metabolites in the small intestine^14^. Thus we hypothesized that in vivo protein tyrosine nitration may occur in *V. cholerae*. Hence, we designed an experiment to check the in-vivo protein nitration pattern of *V. cholerae* cells in a growth phase dependent manner. For this *V. cholerae* (strain C6706) cells grown in LB media were collected at different growth phases and in vivo protein nitration was detected in cell lysates using immunoblotting with monoclonal anti nitrotyrosine antibody. The results showed an extensive protein nitration profile in all the growth phases. Interestingly, the extent of protein nitration was found to be increased during log phase and it reached maxima in the late log phase. Protein nitration was rather low in stationary phase compared to late log phase (Fig. 1). To understand whether this extensive protein nitration pattern is culture media specific or not, we checked the nitration profile of *V. cholerae* cells by growing it in M9 minimal media supplemented with 0.4% glucose. However, the trend of protein nitration profile was almost similar to that of *V.cholerae* cells grown in LB media (Fig. 2). Thus it can be concluded that in vivo protein nitration in *V. cholerae* is not media specific and cells happily grow along with these numerous nitrated proteins.

**Fig. 1.**
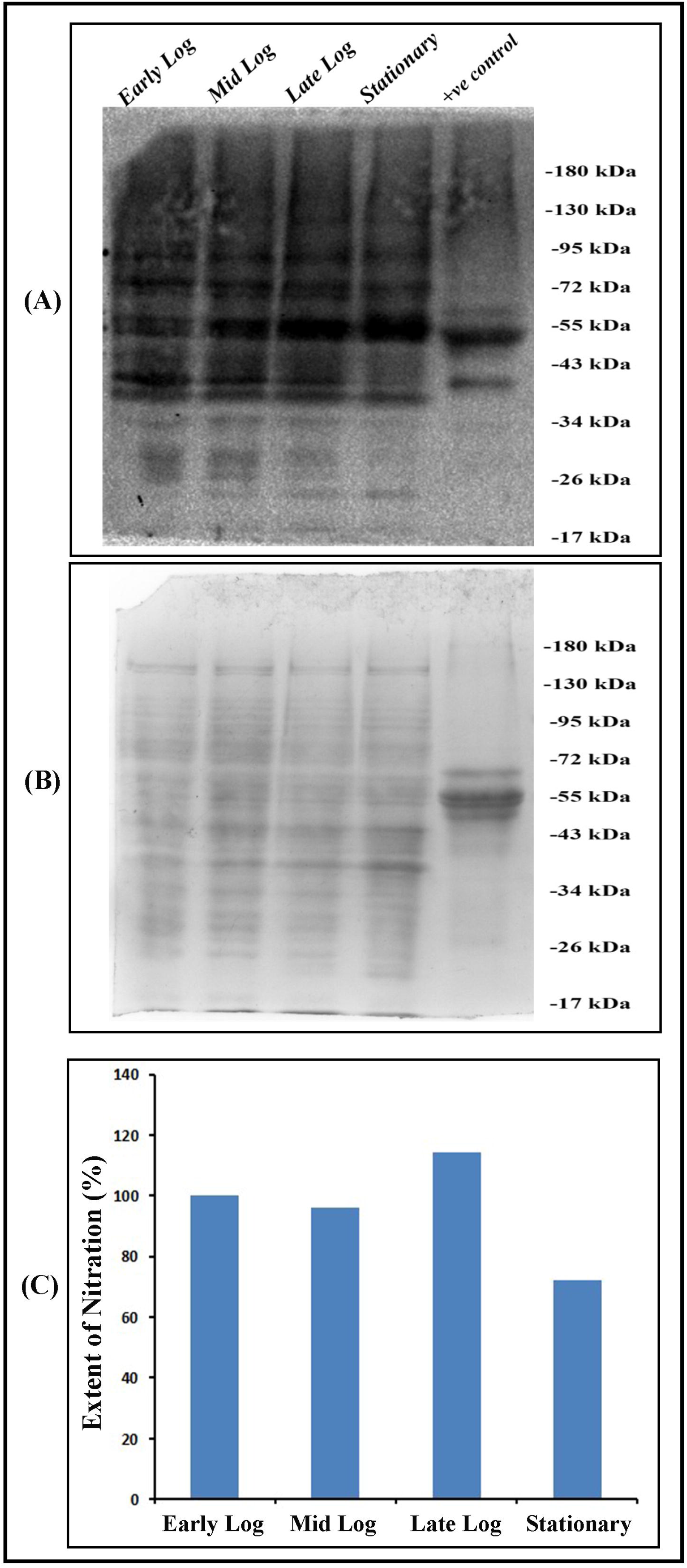
In-vivo growth phase specific Nitration profile of *V. cholerae* (C6706) cell grown in LB media. Lysates from different growth phases of *V. cholerae* cells grown in LB media were subjected to immunobloting using anti 3-nitrotyrosine monoclonal antibody. (A) Western Blot, (B) partially transferred commasie stained gel as loading control and (C) Densitometric analysis of nitration profile from blot.

**Fig. 2.**
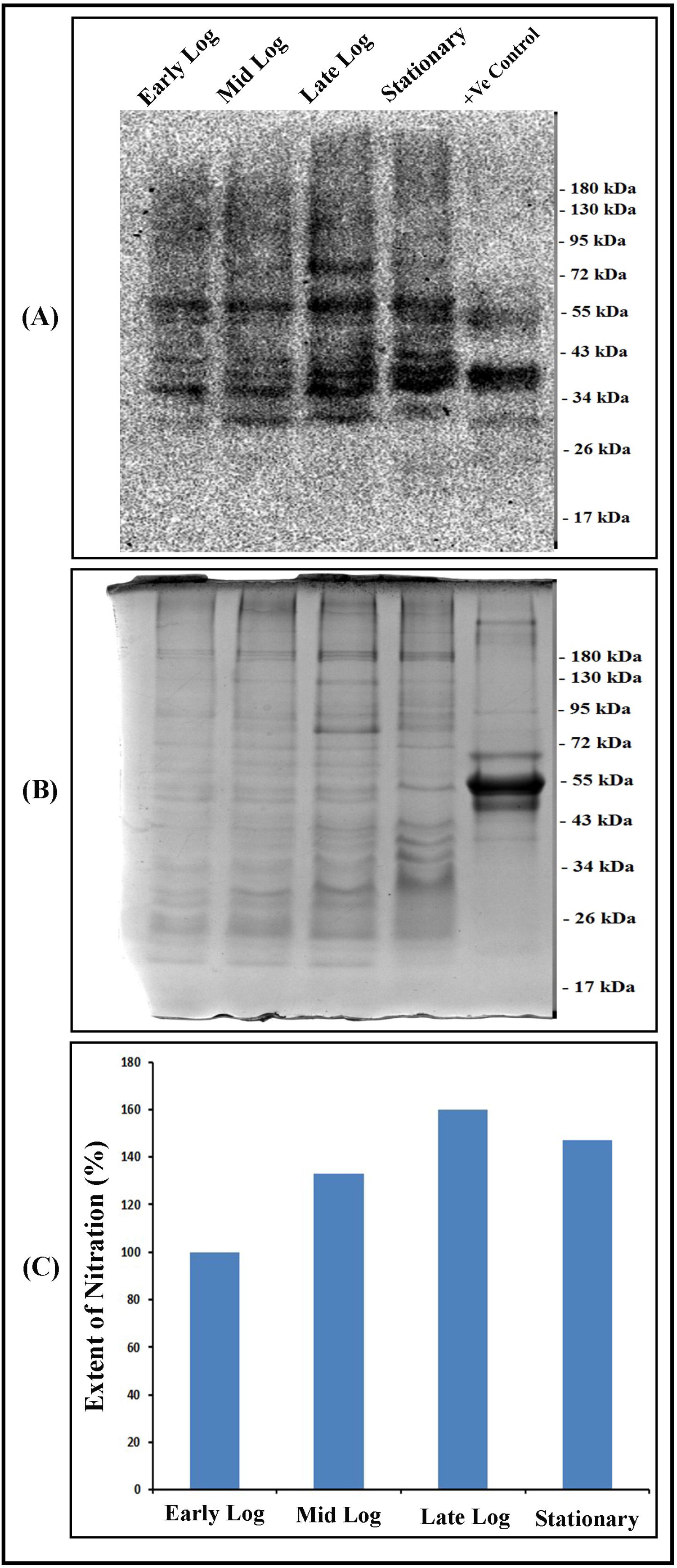
In-vivo growth phase specific Nitration profile of *V. cholerae* (C6706) cell grown in M9 minimal media. Lysates from different growth phases of *V. cholerae* cells grown in M9 media were subjected to immunobloting using anti 3-nitrotyrosine monoclonal antibody. (A) Western Blot, (B) partially transferred commasie stained gel as loading control and (C) Densitometric analysis of nitration profile from blot.

### Identification of nitrated proteins in *V. cholerae* by LC-ESI-MS/MS analysis

Next we tried to identify the nitrated proteins using LC-MS analysis. As we observed numerous nitrated proteins in our previous experiment using the El Tor O1 strain C6706, we tried to confirm the same in another strain N16961 belonging to the same group. For this, *V. cholerae* cells belonging to C6706 and N16961 strains were grown in LB media till the late log phase, where maximum extent of nitration was found. In LC-ESI-MS/MS proteomic analysis, we could identify 29 nitrated proteins from C6706 strain and 58 nitrated proteins from N16961 strain. Interestingly, proteins were nitrated either in single or in multiple tyrosine residues at 3- position of phenyl ring [Table-S1, S2 & S3]. There are several physiologically important proteins and enzymes in the list of nitrated proteins identified from both the O1 serotype strains. Among these proteins, several Glycolytic pathway, TCA cycle and ETC related enzymes are noticeable, i.e Phosphoenol pyruvate carboxykinase (ATP), Pyruvate dehydrogenase E1 component, Fructose-bisphosphate aldolase, Dihydrolipoyl dehydrogenase, Glycerol kinase, Aconitate hydratase B and Enolase etc. Along these we also found several metabolically important proteins like Aldehyde-alcohol dehydrogenase, GDP-mannose 4_6-dehydratase, Glutathione Reductase, Glutamine synthetase, Tryptophanase etc. as nitrated in multiple tyrosine positions. Chaperone protein like 60 kDa chaperonin, ribosomal subunit proteins such as 50s ribosomal L11, L5, L21 are also important nitrated proteins in that list (Fig 3).

**Fig. 3.**
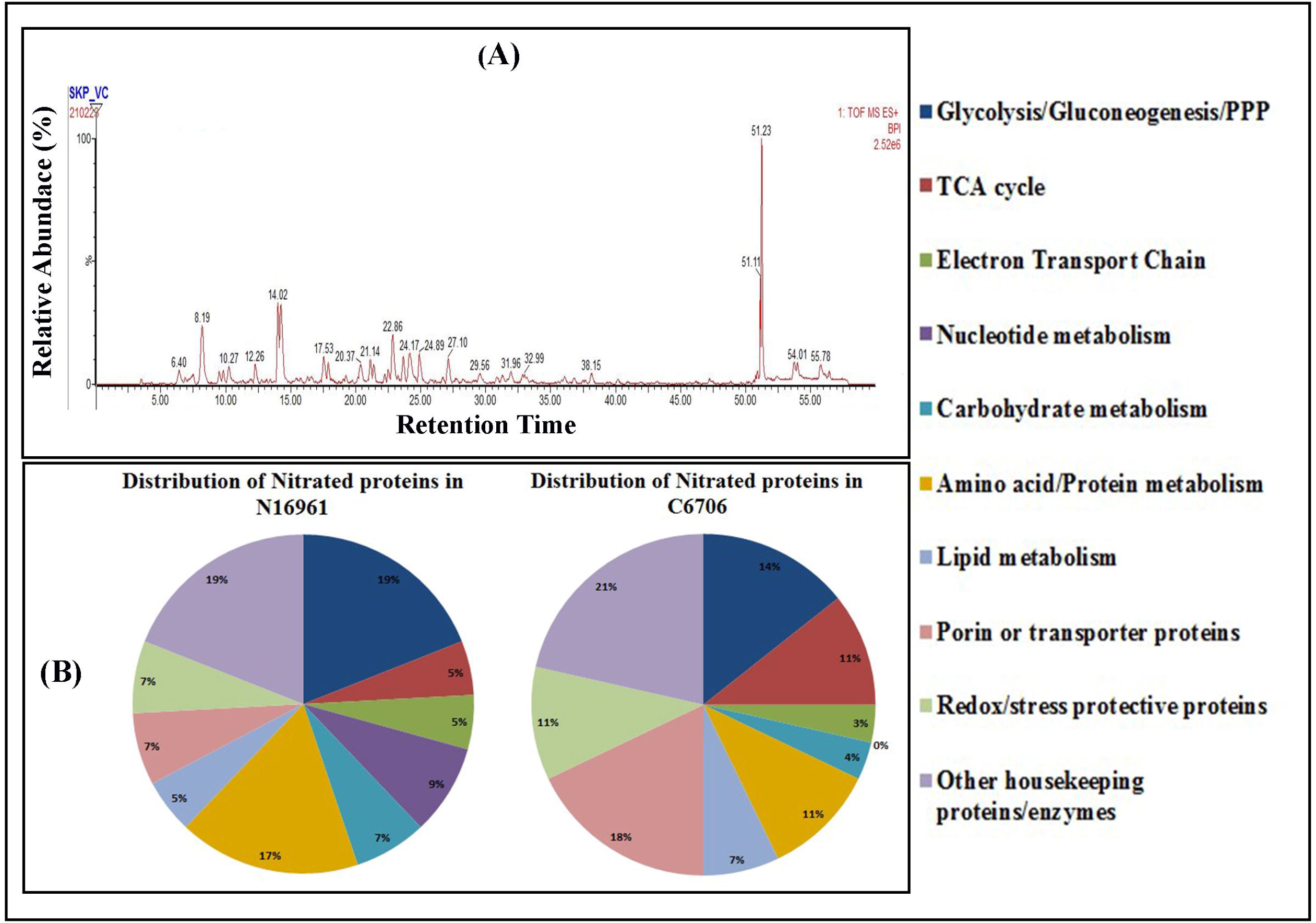
Results of proteomic analysis of *V. cholerae* proteome. (A) Chromatogram of LC-ESI-MS/MS proteomic analysis for the identification of nitrated proteins in *Vibrio cholerae* strain C6706 (B) Pathway specific distribution of identified nitrated proteins of both C6706 and N16961 strains.

### Comparative analysis of intracellular nitrite content of different growth phases of *V. cholerae* grown separately in rich media and in minimal media

It has been reported that protein tyrosine nitration can be mediated by peroxynitrite as well as other species like NO_2_^-^ or •NO_2_. Protein tyrosine nitration can occur through Myeloperoxidase in the presence of NO_2_^-^ and H_2_O_2_ which is an important process in the neutrophil degranulation at the inflammation site. Biological tyrosine nitration can also involve the action of NO_2_^-^ under acidic conditions in the gastric lumen^15^. So along with the nitration profile, we further checked the intracellular nitrite content of *V. cholerae* cell free extracts (CFE) from different growth phase cells grown both in LB and M9 minimal media (Fig.4). Surprisingly, we found a drastic decrease in the intracellular nitrite content as the *V. cholerae* cells progress towards log phase in LB media. Similar result was seen in the *V. cholerae* cells collected from M9 media but the extent of decrease in the intracellular nitrite content was much less, compared to the cells grown in LB. It is important to mention that in both the cases, late log phase cells showed lowest intracellular concentrations of nitrite. Intracellular nitrite content was found to be increased significantly in the stationary phase *V. cholerae* cells collected from both LB and M9 media. Interestingly, we observed a negative correlation pattern between the extent of protein nitration profile and the intracellular nitrite content of *V. cholerae* cells collected at different phases of growth i.e. higher the nitrite content of cells lower is the extent of nitration or vice versa.

**Fig. 4.**
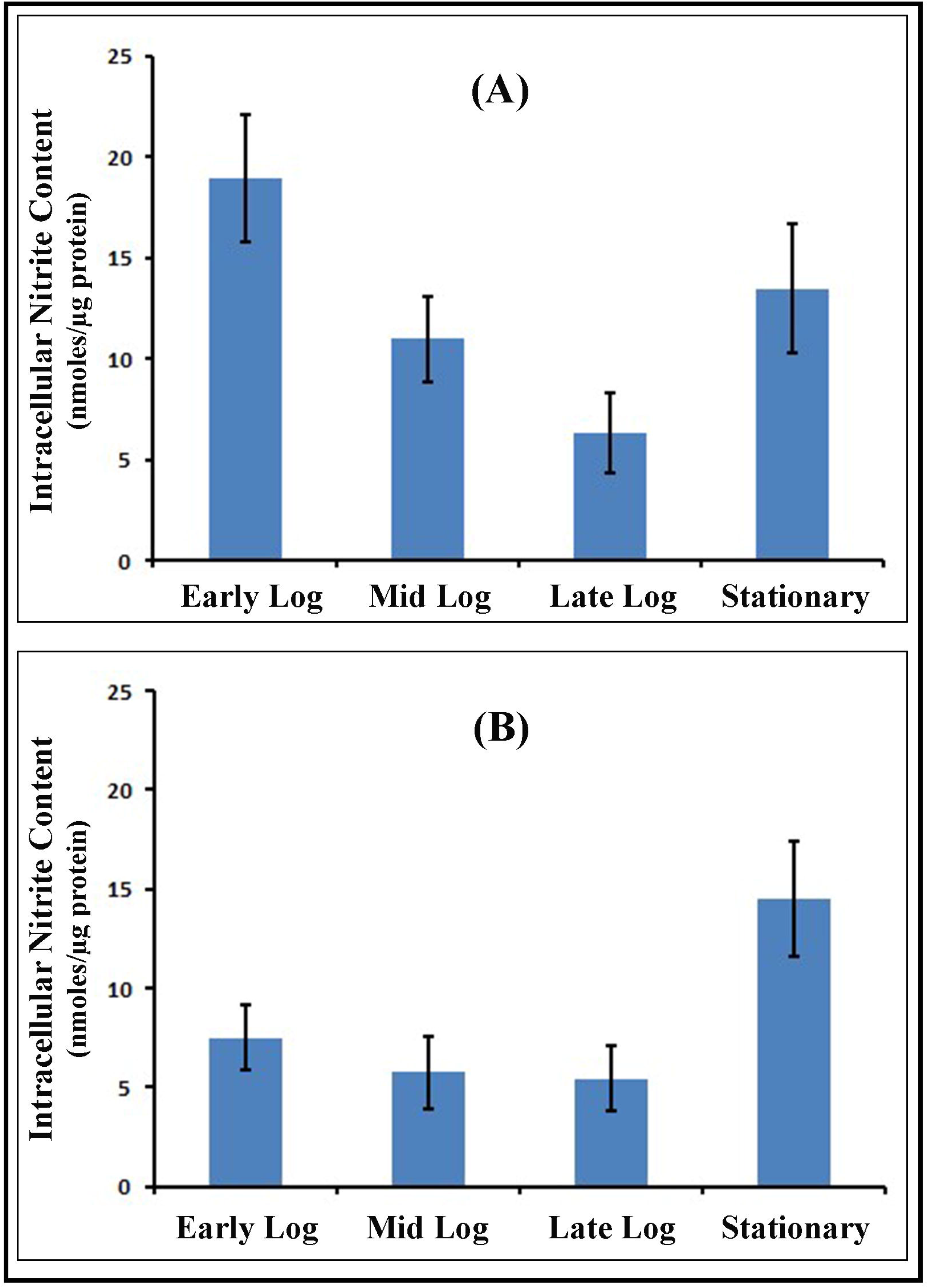
Intracellular Nitrite content of *V. cholerae* cells taken from different growth phase. When, grown in (A) LB media and (B) M9 minimal media. Nitrite content is expressed as concentrations in the terms of nmoles/μg proteins unit. This data is represented as mean ± SD (n = 3).

### Nitrite could induce protein nitration in log phase *V. cholerae* cells grown in minimal media under aerobic condition

Generally, culture media contain nitrate as well as nitrite that can facilitate the phenomenon of nitration in cells. As we found an interesting correlation between intracellular nitrite content and nitration profile in our previous experiments, so we further questioned ourselves whether nitrate or its reduced form nitrite is responsible for this in-vivo nitration. Thus we designed an experiment where we incubated early log phase *V. cholerae* cells grown in M9 minimal media with 1 mM KNO3 and 1 mM NaNO_2_ separately for 2 hours (till mid log phase) along with their untreated control set at 37°C under shaking condition aerobically. After the treatment we prepared cell lysates and checked their nitration profiles using monoclonal anti nitro tyrosine antibody (Fig. 5A). Densitometric analysis of the protein nitration blot showed a significant increase (~ 57%) in nitration in 1 mM nitrite treated early log phase *V. cholerae* cells compared to control set grown under similar experimental conditions. But no such significant increase (~12%) in extent of protein nitration was observed in 1 mM nitrate treated *V. cholerae* cells (Fig. 5C). So, these results directly point toward the clear fact of nitrite mediated in-vivo PTN in *V. cholerae*.

**Fig. 5.**
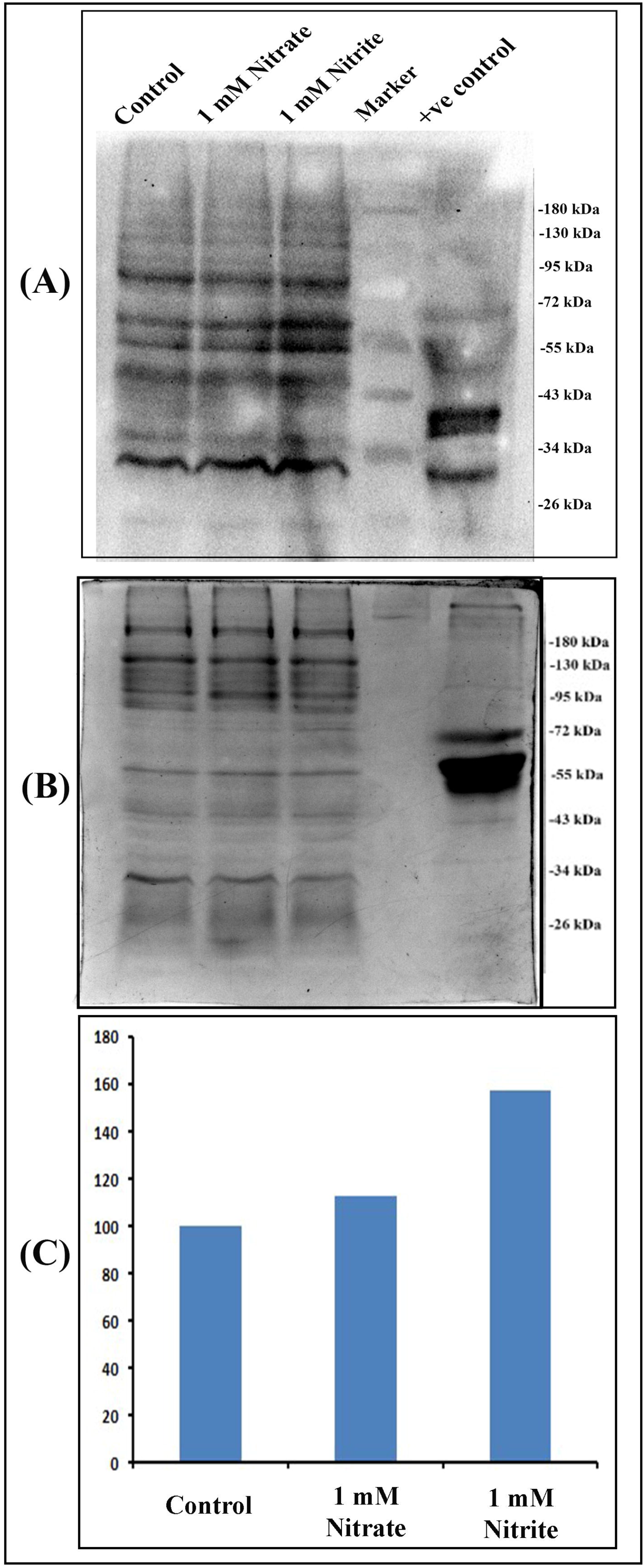
Nitration profile of *V. cholerae* proteome after treatment with 1 mM nitrate and 1 mM nitrite. *V. cholerae* was grown for 2 hours in M9 minimal media in the presence of 1 mM nitrate and 1mM nitrite during early log phase growth and the nitration profile was checked. (A) Western blot, (B) partially transferred commasie stained gel as loading control and (C) Densitometric analysis of nitration profile from blot.

### Protein denitration was induced in *V. cholerae* cells under anaerobic conditions

PTN is generally considered as a stress induced post translational modification in biology, which causes mostly altered functions of proteins. But here we observed that PTN is a natural phenomenon occurring in-vivo during aerobic growth phase of *V. cholerae* cells. To evaluate the status of protein nitration found in aerobic growth, we designed an experiment where, we did an en masse transfer of late log phase *V. cholerae* culture grown in aerobic condition into anaerobic condition and incubated that system at 37°C in shaking condition for 18 hours along with its aerobic set. Then we checked the protein nitration profile of anaerobically grown cells against its aerobic stationary control set using western blot analysis. Our results showed a significant 37% decrease in the extent of nitration of the anaerobic set compared to its aerobic control set (Fig. 6). Thus it can be concluded that *V. cholerae* cells showed in-vivo protein tyrosine nitration during aerobic growth and protein de-nitration was prominent during anaerobic or hypoxic phase.

**Fig. 6.**
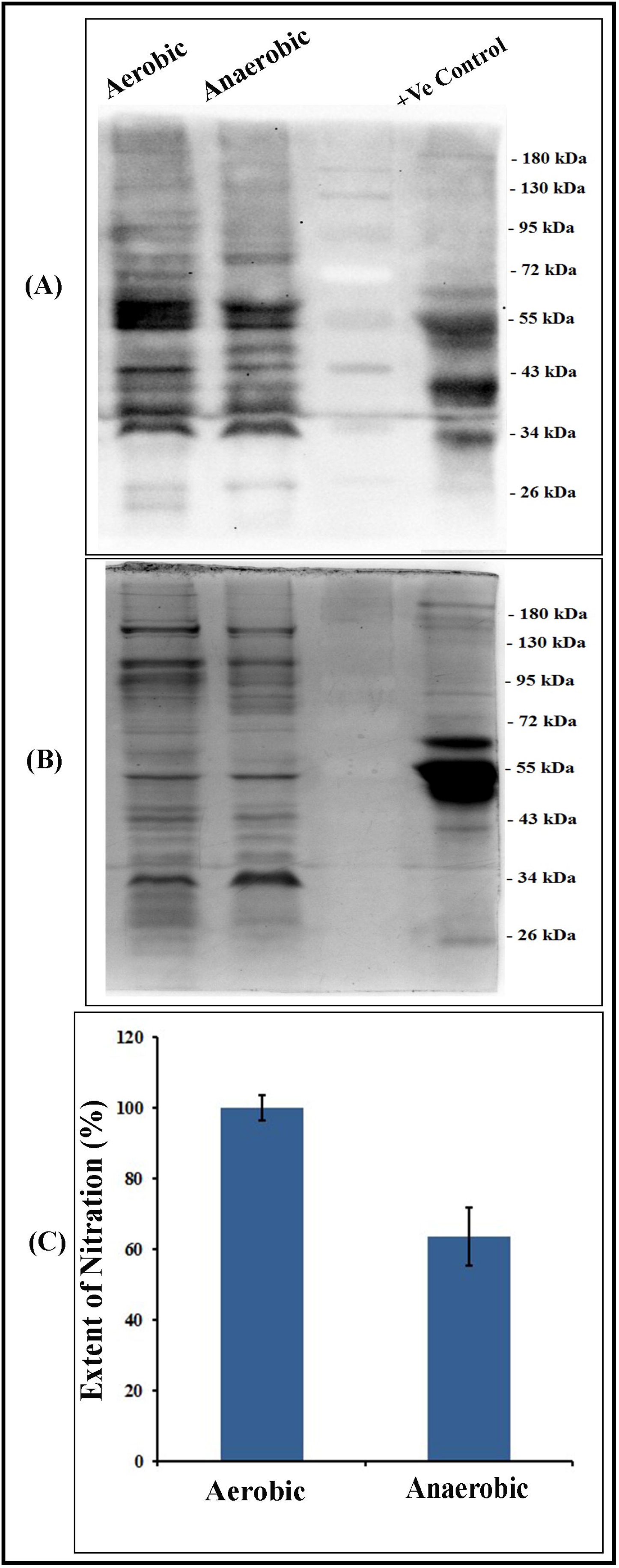
In-vivo nitration profile of anaerobically incubated *V. cholerae* proteome. Aerobically grown late log phase *V. cholerae* cells were subjected to anaerobic incubation for 18 hours at 37°C under shaking condition along with its aerobic control set in M9 minimal media. (A) Western blot, (B) partially transferred commasie stained gel as loading control and (C) Densitometric analysis of nitration profile from blot.

### Nitrite to nitrate conversion facilitate stationery phase *V. cholerae* cell survival under anaerobic condition

It has been reported in earlier studies that nitrate reductase enzyme is essentially required for nitrate dependent anaerobic respiration in different microorganisms as well as in *V. cholerae*, where the enzymatic reduction product of nitrate is nitrite^16,17^. In our previous experiments, we found a direct evidence of nitrite mediated protein nitration in *V. cholerae* cells and a denitration phenomenon was also observed in *V. cholerae* proteome during anaerobic incubation at stationary phase. So, keeping these findings in mind, we checked the intracellular nitrite as well as the nitrate content of *V. cholerae* stationary phase cells incubated in anaerobic conditions and compared them with aerobically incubated cells. We found no significant increase in nitrite content of anaerobically incubated stationary phase cells (5 ± 1.7 nmoles of NO_2_/μg proteins) compared to its aerobic control set (3.8 ± 0.37 nmoles of NO_2_/μg proteins). But we found a significant increase (~ 2.2 fold) in nitrate in anaerobic set compared to aerobic control set. The concentration of nitrate in aerobically grown *V. cholerae* was 2.78 ± 0.3 nmoles nitrate /μg protein whereas 6.2 ± 1.1 nmoles nitrate/μg protein was found in anaerobic set up (Fig. 7). This result is corroborated well with previous report where exogenous nitrate treatment under anaerobic condition facilitated *V. cholerae* cell survival^18^. Thus it can be concluded that cellular nitrate pool is enriched under anaerobic condition which actually facilitate cell survival under anaerobic respiration. Moreover, increased nitrate content is directly linked with the de-nitration of proteome which ensures supply of nitrite into intracellular environment.

**Fig. 7.**
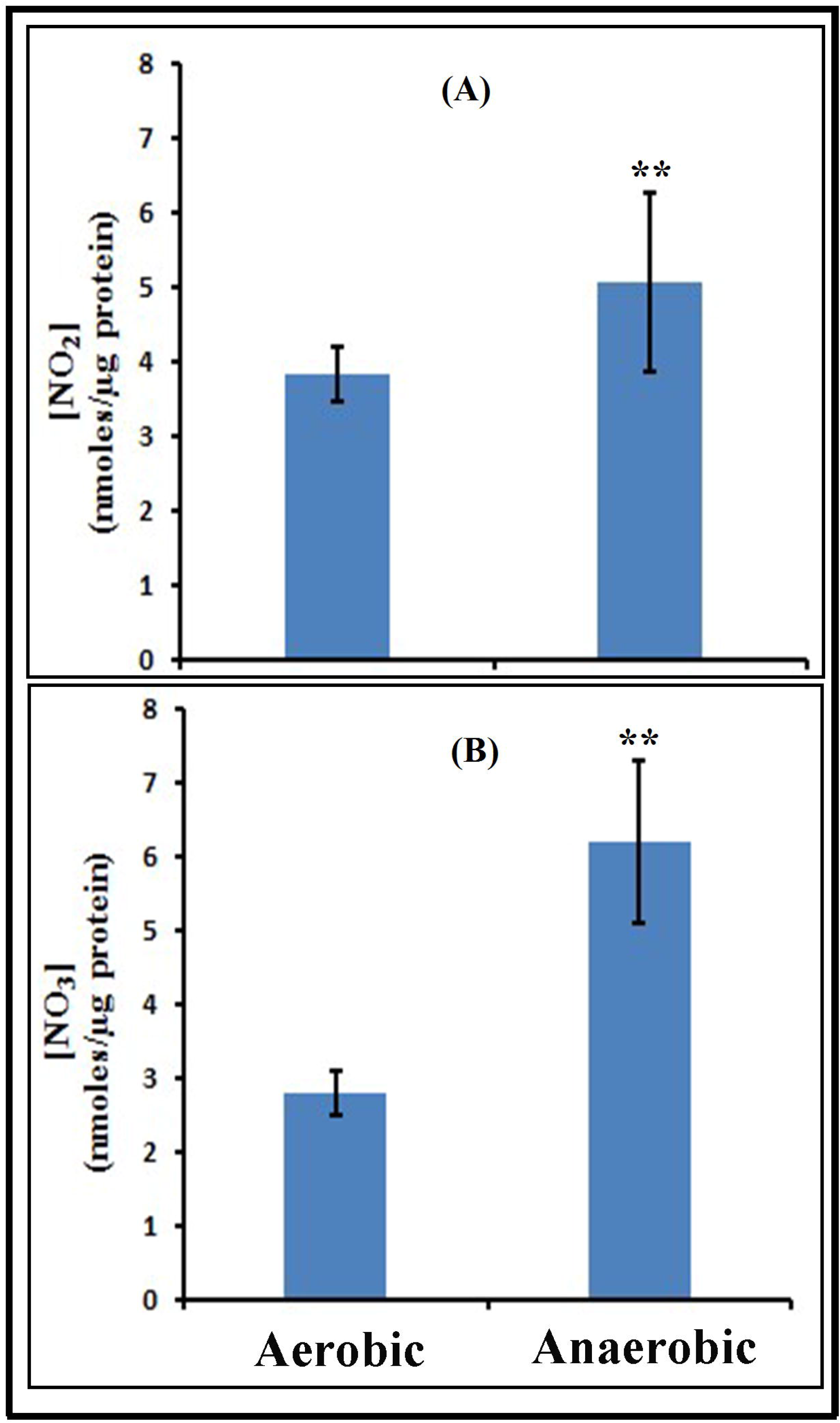
Determination of intracellular nitrite and nitrate content of anaerobically incubated *V. cholerae* cells. *V. cholerae* cells were incubated for 18 hours anaerobically in shaking condition at 37°C in M9 minimal media. (A) Intracellular nitrite and (B) intracellular nitrate content were measured in aerobically and anaerobically incubated late log phase cells. Nitrite and nitrate content is expressed as concentration with nmoles/μg protein unit. This data is represented as mean ± SD (n = 3). **p < 0.01.

### Cellular redox status supports nitrite to nitrate conversion under anaerobic condition

In our previous experiment we found an increased intracellular nitrate content in anaerobically incubated stationary phase cells compared to its aerobic control set. Protein denitration generates soluble nitrite inside the cell which is not useful for the *V. cholerae* but also highly toxic. So the nitrite generated from the de-nitration must have been oxidized and converted into the nitrate inside the cells to keep a continuous supply of nitrate for cell survival under anaerobic condition. Intracellular environment is reducing in nature in all types of cells starting from prokaryotes to higher eukaryotes except in Golgi apparatus where oxidizing environment is required. So, to check the possibility of nitrite to nitrate conversion inside the stationary phase *V. cholerae* cells, we assessed the redox environment of the same under anaerobic as well as aerobic conditions.

We determined total glutathione, oxidized glutathione (GSSG), reduced glutathione (GSH), as well as GSH/GSSG ratio. The GSH content was significantly higher in aerobic set of *V. cholerae* cells (46.35 ± 4.5 nmole/mg protein) than the anaerobic set (29.3 ± 7 nmole/mg protein). However, the GSSG content was not increased much in anaerobic set (2.58 ± 0.2 nmole/mg protein) compared to its aerobic counterpart (1.2 ± 0.26 nmole /mg protein). Thus the significant decrease in GSH content of stationary phase *V. cholerae* cells incubated under anaerobic conditions ultimately resulted in ~3.5 fold decrease in GSH/GSSH ratio compared to its control aerobic set (Fig. 8). So it can be concluded that a relatively oxidizing environment is prevailed inside *V. cholerae* cells under anaerobic condition, which promote the conversion of nitrite to nitrate and facilitates the cell survival.

**Fig. 8.**
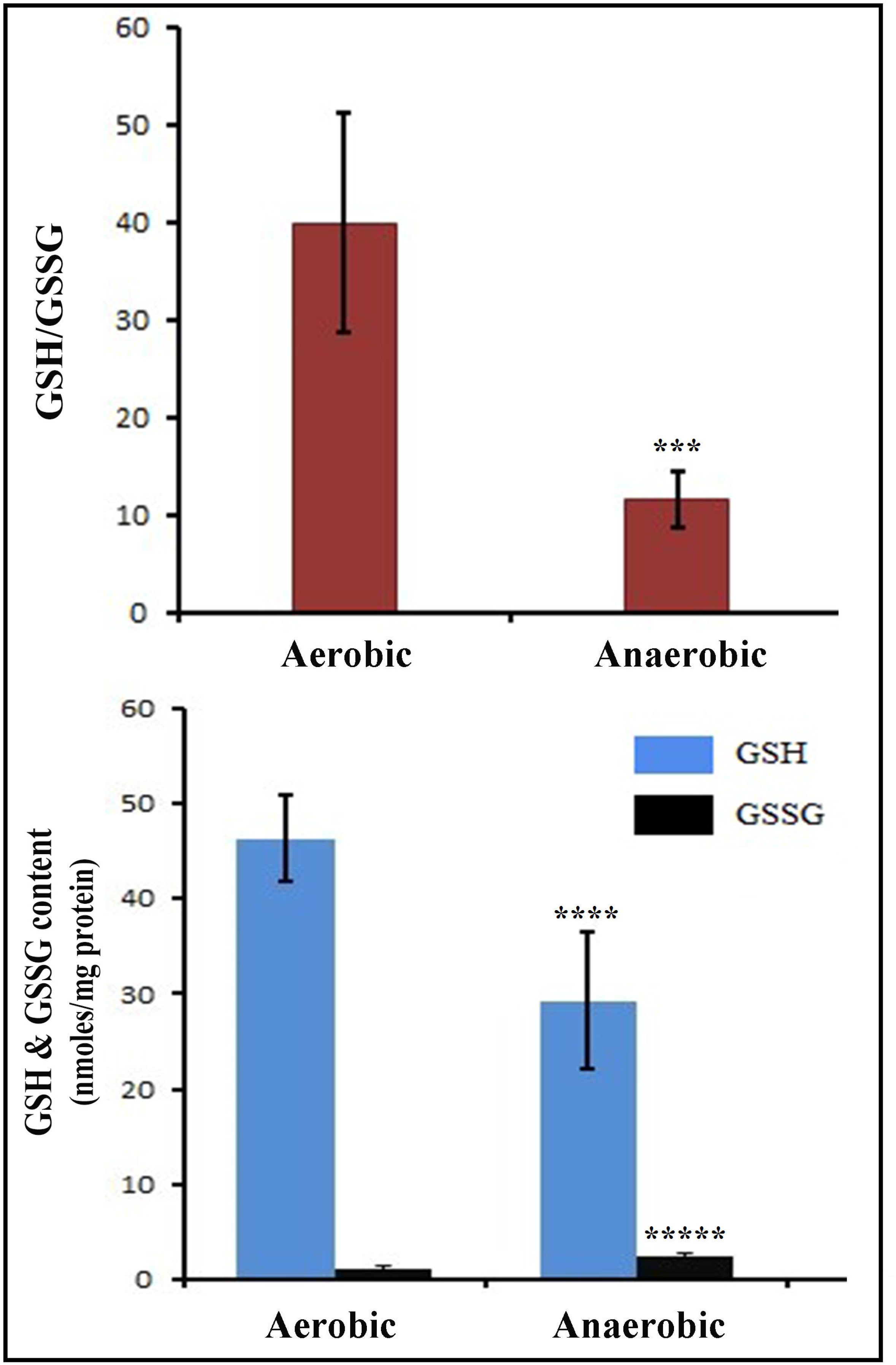
Determination of cellular redox status of *V. cholerae* incubated under aerobic and anaerobic condition. Cellular redox status was assessed in aerobically and anaerobically incubated late log phase *V. cholerae* cells which were grown for 18 hours in shaking condition at 37°C in M9 minimal media. Redox status represents the reduced glutathione (GSH) concentration, oxidized glutathione (GSSG) concentration and GSH/GSSG ratio. Where, GSH and GSSG concentrations are expressed in nmoles/mg protein unit. This data is represented as mean ± SD (n = 3). ***p < 0.001, ****p<.0001.

## Discussion

The virulence factors and the pathobiology of the disease cholera are well characterized^19^. Inside the human gut, *V. cholerae* faces several host inflammatory stresses including nitrosative and oxidative stress. In this stressful condition post translational modification like protein tyrosine nitration (PTN) and its related detrimental effect is quite eminent as well as unavoidable by any gut pathogen. Along with this, presence of a prominent hypoxic environment in gut should make *V. cholerae* hard to survive theoretically. But the practical scenario is quite different which shows that *V. cholerae* cells are quite capable of coping up with these harsh conditions not only by continuing the survival but also by population expansion.

In our study we found in-vivo protein tyrosine nitration in *V. cholerae* irrespective of particular growth media. Interestingly, the extent of nitration was found to be highest during its log phase. This indicates that unlike most organisms where protein nitration is overall detrimental, *V. cholerae* is not only tolerant to PTN but it can naturally thrive with this post translational modification on multiple proteins. Most interestingly, our LC-ESI-MS/MS data showed that these multiple proteins include housekeeping proteins as well as important metabolic proteins. In the search of the source, we found that in-vivo nitration is directly governed by nitrite. During our study we came across a very in-depth research article related to *V. cholerae* published by *Bueno et al*.,^18^ in the year 2018. *Bueno et al*. gave an elaborate relation among the following: i) nitrate reductase (napA) meditaed anaerobic respiration, ii) low pH dependent increased cell viability with decreased population expansion and iii) exogenous nitrate treatment. They found higher level of nitrite in growth media when they administrated exogenous nitrate during hypoxic growth, as *V. cholerae* lacks nitrite reductase^20^. In addition to their study, our finding of in-vivo nitration in aerobic culture showed another possible fate of endogenous nitrite. In-vivo nitration could also be another way to entrap more toxic nitrite without causing harm to cell in anaerobic condition. In the quest to understand the significance of in-vivo nitration, we further found de-nitration phenomenon of *V. cholerae* proteome and increase in intracellular nitrate during anaerobic incubation of aerobically grown late log phase culture. In a similar experiment when we imparted anaerobicity to *V. cholerae* culture grown aerobically till early log phase, we found reverse results i.e. increased in-vivo nitration in anaerobic set (Supplementary Fig. 1). The different observations are due to the nutrient i.e. nitrate rich conditions of mid log culture media compared to nutrient deficient i.e. nitrate deficient late log phase media. This result also indirectly proved our theory of nitrite induced in-vivo nitration because of *napA* overexpression during anaerobic condition. This made us to understand that in-vivo nitration might help *V. cholerae* cells to keep a reservoir of toxic nitrite in a “safe condition” for future use. This denitration phenomenon is synergistically associated with low GSH/GSSH of cell i.e. more oxidative environment that is prevailed during long term anaerobic incubation, where released nitrite is oxidized and converted to nitrate which can be used during anaerobic respiration. Hence in-vivo nitration phenomenon is important for *V. cholerae* cell survival mostly in a condition where scarcity of exogenous nitrate is prevailing. From these facts, we propose that in a nitrate deprived anaerobic growth condition, in-vivo nitration and de-nitration is a cyclical phenomenon which is governed by controlled oxidative condition of cell and can perform as a sole source of respiration just to keep cells viable. Apart from this, in our previous studies we showed the role of S-nitrosoglutathione Reductase (GSNOR)^21^ and Catalases (KatB & KatG)^22^ in combating nitrosative stress mediated by GSNO and peroxynitrite respectively. Hence, in-vivo nitration might also be an advanced evolutionary adapted mechanism of nitrosative stress tolerance found in *V. cholerae*, as it restricts proteome to unnecessary PTN in nitroso-oxidative stress prevailing environment such as inside human gut.

Perhaps there are many possibilities of in-vivo nitration in other organisms, which face similar harsh environment as *Vibrio sp.*, or in those organisms which have alternate habitats consisting of aerobic and anaerobic environments. Even in our study we found lowered extent of nitration and higher intracellular nitrite content in stationary phase cells. This result might indicate the possible role of de-nitration as well as the role of *napA* in cell survival during nutrient deprived state.

## Methods

### Strains and media used

In this study *Vibrio cholerae* strain C6706 belonging to El Tor O1 serotype was used throughout all experiments. For growth purpose, LB media with 1% NaCl (pH 7.5) and M9 minimal media with 0.4% glucose (pH 7.5) were used. For proteomic analysis (LC-ESI-MS/MS) N16961 belonging to El Tor O1 was used along with C6706.

### Aerobic and anaerobic growth conditions for nitration/de-nitration studies

For aerobic growth conditions cells were grown in either LB or M9 minimal media with a starting O.D.600nm of 0.05 in shaking condition (140 rpm) at 37°C. The growth was monitored by checking the O.D.600nm at 1 hour to 2 hours interval.

To impart anaerobicity, cells were grown aerobically till late log phase, then, the whole culture media containing cells were transferred to closed screw capped tubes with negligible void volume (air) and grown for further 18 hours at 37°C in shaking condition to study stationary phase denitration.

### Preparation of Cell free extract (CFE) for immunoblotting and other assays

Cells were collected by centrifugation at 5000g for 10 minutes followed by washing the cell pellet with PBS and recollecting it by centrifugation. Collected cells were resuspended and lysed with 20 mM Tris-HCl buffer of pH 7.5, containing 1 mM EDTA, 1 mM PMSF and 0.5 mg/ml lysozyme followed by incubation at 37°C for 10 minutes and finally by sonication. The CFE is finally collected by centrifugation of lysate at 10000g for 10 minutes. Protein concentrations of respective CFE samples were checked spectrophotometrically by using Bradford reagent at 595nm.

### Detection of protein tyrosine nitration (PTN) by western blot analysis

To detect nitration in whole proteome, equal and normalized amount of protein (30μg) samples were prepared without adding β-mercaptoethanol (BME) and were subjected for SDS-PAGE using 5% stacking and 10% resolving gel of 1.5 mm width. After the electrophoresis process, proteins within the gel were partially transferred to polyvinylidene difluoride (PVDF) membrane (250 mA current flow, 1.5 h) using wet transfer apparatus (Bio-Rad Laboratories Inc., Hercules, CA, USA). Blocking of the PVDF membranes were done for overnight at 4°C temperature in shaking condition using blocking buffer (0.019 M Tris, 0.136 M NaCl, 0.1% v/v Tween 20 and 5% w/v nonfat dry milk). Anti 3-nitrotyrosine primary antibodies at 1:2000 dilutions in tris-buffered saline with Tween 20 (TBST) (0.019 M Tris, 0.136 M NaCl, 0.1% v/v Tween 20) were used to probe the membranes at room temperature for 1.5 hour. The primary antibody probed membranes were washed three times in TBST for 10 min for each wash, and then re-probed with HRP-conjugated anti-mouse IgG antibody at 1:2000 dilutions for 1 h at room temperature. The membrane was then finally washed three times in TBST followed by three times in TBS (0.019 M Tris, 0.136 M NaCl) (10 min for each wash). The immunopositive spots were visualized by using chemiluminescent reagent (Thermo Scientific Pierce, Rockford, IL, USA) as directed by manufacturer. The extent of relative nitration was measured by densitometric scanning of the immunopositive blot using ImageJ software.

### Detection of protein tyrosine nitration and identification of nitrated proteins using LC-ESI-MS/MS Proteomic analysis

*Vibrio cholerae*, C6706 strain was used for proteomic analysis study. Cell lysates of mid log phase cells grown in LB media were prepared in the same way previously described in cell lysate preparation column. The cell lysates were first lyophilized and then dissolved in 50mM ammonium bicarbonate solution such a way that all protein samples would reach to same concentration (2 mg/ml protein). Crude cell lysates were then subjected to treatment by TFE followed by reduction by 5mM DTT for 1 hour at 60°C. Then alkylation of sample was done by 5mM Iodoacetamide at room temperature for 45 minutes. After these, sample was subjected to overnight trypsin digestion. Trypsin digested samples were treated with 0.1% formic acid and subjected for Liquid chromatography (LC) followed by ESI-MS/MS detection and analysis in XEVO G2-XS QTof system (Waters). BEH C-18 column (Waters) was used in the instrument. Sodium formamide was used as primary standard and whereas, leucine encaphline was used as secondary standard. Ramp collision energy was set at 18 to 40 V. whereas; MS and MS/MS thresholds were set at 150 and 20 counts respectively. Only fresh cell lysates were used in this study to get the best possible quantitative results. Protein tyrosine modifications in peptides were determined by setting a parameter indicating with mass shift of +47Da^23^. The peptides detected by LC-MS/MS were matched with Vibrio cholerae, C6706 database downloaded from Uniprot in FASTA format. The software used for analysis was Progenesis Qip by Waters.

### Determination of intracellular nitrite (NO_2_) concentration

Intracellular nitrite content was determined using Griess assay^24^ in freshly prepared CFE. Briefly, CFE was incubated at room temperature in dark with 1% Sulfanilamide solution prepared in 5% phosphoric acid for 15 minutes followed by addition of equal volume of 0.1% N-(1-naphthyl) ethylenediamine dihydrochloride (NEDD) solution and incubation for 15 minutes in similar way. The resulting azo dye formation inside sample tubes was spectrophotometrically measured at 540 nm against a substrate blank. The concentration of nitrite was determined from the O.D. value using a standard curve of nitrite made using same reagents and experimental setup.

### Treatment of *V. cholerae* cells with NO_3_ and NO_2_

To see the pattern of protein nitration in *V. cholerae* in response to exposure of nitrate and nitrite, *V. cholerae* cells were grown and incubated aerobically in separate conical flasks with 1mM potassium Nitrate (KNO_3_) and 1 mM Sodium Nitrite (NaNO_2_) from lag phase to mid log phase in M9 minimal media along with an untreated control set. To observe cellular growth, culture O.D. was monitored spectrophotometrically at 600 nm in a 1-hour regular interval until the O.D reached around 0.8 starting from around 0.05. After that the cells were collected and subjected to prepare CFE for immunoblotting.

### Determination of intracellular nitrate (NO_3_) concentration

Intracellular nitrate content was measured by a multi-step process. In brief, at first cellular NO3 was converted to NO_2_ enzymatically using Nitrate Reductase (NR) enzyme followed by few steps, then the cellular nitrite was measured using Griess assay as mentioned earlier. To convert nitrates to nitrites freshly prepared CFE samples in phosphate buffer (pH 7.5) were incubated with 0.1 U/ml NR, 5μM FAD, 30μM NADPH for 30 minutes at 37°C. Then the sample was incubated again for 30 minutes at 37°C with 0.1 kU/ml Lactate dehydrogenase (LDH) and 300μM Pyruvate to ensure full oxidation of NADPH to avoid any interference in Griess reaction. After these steps the samples were subjected to Griess assay and the nitrite content was measured. To measure the nitrate content, this experiment was done with two setups for each sample. One of which is marked as NR blank and that was used to subtract the true nitrite content of samples.

### Total Glutathione (GSH+GSSG), Reduced Glutathione (GSH), and Oxidized Glutathione (GSSG) Content Measurement

These parameters were measured according to the method described by Akerboom et al.^25^. In brief, freshly prepared, crude cell free extract was added to equal volume of 2M HClO4 containing, 2mM EDTA and incubated on ice for 10 min. The mixture was centrifuged at 5000×g for 5 min. Resulting supernatant was neutralized with 2M KOH containing 0.3M HEPES. After centrifugation at 5000×g for 5 min, the neutralized supernatant was used for estimation of the above-mentioned parameters. Total glutathione was estimated following Glutathione Reductase (GR)-dependent DTNB reduction assay spectrophotometrically measuring 5-thio-2-nitrobenzoate (TNB) formation at 412 nm. The reaction mixture contained 100mM K-phosphate, pH 7.0, 0.2mM NADPH, 0.12 U GR, 1mM EDTA, 0.063mM DTNB and sample in a total volume of 500μl. Same neutralized extract was treated with 2-vinylpyridine (50:1, v/v) for 1 h at room temperature. Then, it was used for GSSG estimation using the above method. Reduced Glutathione content was determined from the difference between the total and oxidized glutathione content of the sample.

### Statistical analysis

All results of biochemical assay were expressed as mean ± SD, for n=3. The statistical evaluation was performed with either one way or with two-way ANOVA followed by two tailed paired Student’s t-test; p value≤0.05 or 0.01 was considered significant.

## Acknowledgements

The authors would like to acknowledge Prof. Jun Zhu, Department of Microbiology, Nanjing Agricultural University, Nanjing, Jiangsu, China and Dr. Rupak Kumar Bhadra, Scientist, IICB, Jadavpur, Kolkata, India for providing *V. cholerae* strains used in this study. The authors also thank the Director, Bose Institute, Kolkata for providing the Proteomics Facility. The authors also thank Mr. Souvik Roy, Proteomics Facility, Bose Institute, Kolkata for his excellent technical support.

## Author contributions

S.K.P. and S.G. designed the experiments. S.K.P., N.S., S.C., A.S. and S.R. designed the data collection process and data collection. S.K.P., N.S. and S.R. performed the proteomic analysis. S.K.P., S.C., A.S. and performed the western blot analysis. S.K.P. performed the experiments related to cell growth, nitrate, nitrite measurement and GSH, GSSG estimation. S.K.P. and S.G. analyzed the data. S.K.P. wrote the first draft of the manuscript. S.G. contributed to checking and revising the manuscript.

---

**Supplementary Figure 1.**
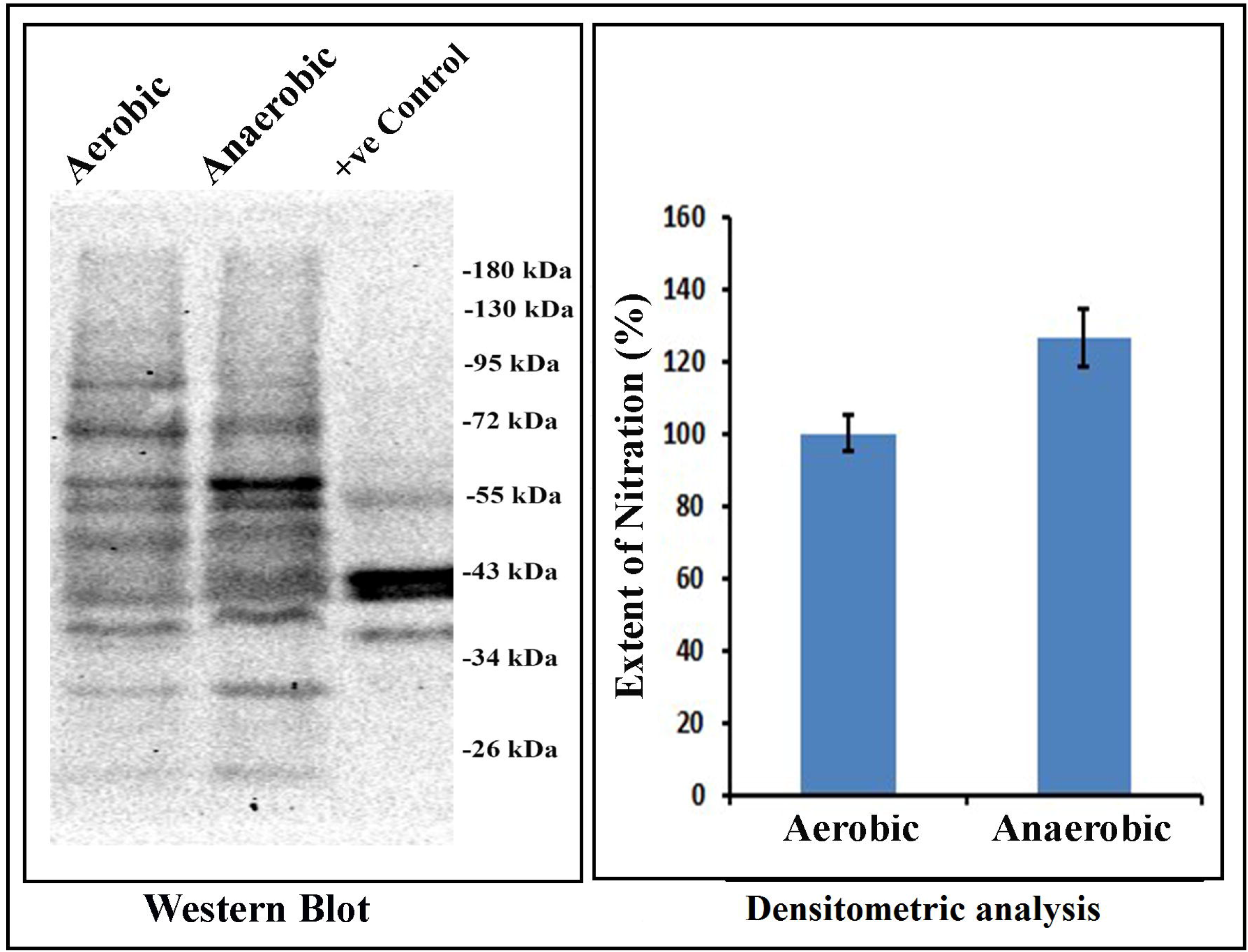

