## Supplementary Table 1 for "In-vivo protein nitration and de-nitration facilitate *Vibrio cholerae* cell survival under anaerobic condition: Consequences of Nitrite induced protein nitration"

**Table S1.** List of identified Nitrated proteins in *V. cholerae* strain N16961

| **Accession No.** | **Description** | **Average Normalized Abundances** | **Score** | **Nitration position** |
| --- | --- | --- | --- | --- |
| [**Q9KQB1**](https://www.uniprot.org/uniprot/Q9KQB1) | **Succinate dehydrogenase flavoprotein subunit** | **1.22e+005** | **41.52** | **Y577** |
| **Q9KSS4** | **Thioredoxin Reductase** | **1.34e+005** | **30.94** | **Y119** |
| **Q9KNH5** | **ATP synthase subunit beta** | **2.33e+006** | **271.91** | **Y275, Y443, Y453** |
| **Q9KPM5** | **Elongation factor G 2** | **1.22e+007** | **503.07** | **Y480** |
| **Q9KUT0** | **50S ribosomal protein L21** | **8.60e+005** | **47.59** | **Y2** |
| **Q9KP11** | **Peptidyl-prolyl cis-trans isomerase** | **3.86e+005** | **46.32** | **Y130** |
| **Q9KSF9** | **Asparagine--tRNA ligase** | **5.55e+005** | **116.70** | **Y3, Y227, Y426** |
| **Q9KV22** | **2_3-bisphosphoglycerate-independent phosphoglycerate mutase** | **4.69e+005** | **97.11** | **Y17, Y366** |
| **Q9KP17** | **2`_3`-cyclic-nucleotide 2`-phosphodiesterase** | **3.93e+005** | **87.60** | **Y521** |
| **Q9KQ29** | **2-dehydro-3-deoxyphosphooctonate aldolase** | **3.27e+005** | **74.40** | **Y41** |
| **Q9KPW8** | **Acetyl-coA carboxylase carboxyl transferase subunit alpha** | **1.99e+006** | **109.99** | **Y140** |
| **Q9KPF5** | **Acetyltransferase component of pyruvate dehydrogenase complex** | **2.91e+005** | **73.77** | **Y606** |
| **Q9KTB7** | **Adenylate kinase** | **6.50e+005** | **79.82** | **Y171, Y181, Y182** |
| **Q9KQG5** | **Aldehyde-alcohol dehydrogenase** | **3.15e+006** | **350.74** | **Y684, Y689, Y774** |
| **Q9KNF1** | **Alpha-1_4 glucan phosphorylase** | **1.11e+006** | **152.48** | **Y316, Y388, Y746, Y756,** |
| **Q9KNN3** | **Aspartate ammonia-lyase** | **4.10e+006** | **267.29** | **Y85, Y158, Y167, Y191** |
| **Q9KNH3** | **ATP synthase subunit alpha** | **5.87e+006** | **155.07** | **Y308,** |
| **Q9KQA8** | **Citrate synthase** | **2.31e+006** | **167.53** | **Y74, Y85, Y94, Y181, Y311** |
| **Q9KPC4** | **CTP synthase** | **4.26e+005** | **133.61** | **Y298** |
| **Q9KPF6** | **Dihydrolipoyl dehydrogenase** | **2.05e+006** | **160.45** | **Y341, Y352** |
| **Q9KV30** | **DNA-directed RNA polymerase subunit beta (rpoB)** | **7.36e+006** | **565.61** | **Y262, Y1228** |
| **Q9KPC5** | **Enolase** | **5.44e+006** | **278.20** | **Y202, Y304** |
| **Q9KQY1** | **Formate acetyltransferase** | **9.96e+006** | **473.85** | **Y116, Y215** |
| **Q06952** | **GDP-mannose 4_6-dehydratase** | **1.95e+006** | **180.48** | **Y85, Y158, Y164, Y166** |
| **Q9KRB5** | **Glucose-1-phosphate adenylyltransferase 1** | **2.93e+005** | **45.47** | **Y239** |
| **Q9KNJ2** | **Glutamine synthetase** | **3.62e+006** | **290.34** | **Y327, Y335, Y288, Y297, Y298** |
| **Q9KVG0** | **Glutathione reductase** | **1.01e+006** | **57.82** | **Y11, Y384** |
| **Q9KQJ8** | **Glyceraldehyde-3-phosphate dehydrogenase** | **6.89e+006** | **413.49** | **Y47** |
| **Q9KLJ9** | **Glycerol kinase** | **5.20e+006** | **361.46** | **Y332, Y344, Y356** |
| **Q9KRC1** | **HTH-type transcriptional repressor PurR** | **6.33e+004** | **17.50** | **Y45** |
| **Q9KVA0** | **Iron-containing alcohol dehydrogenase family protein RfbM** | **9.81e+005** | **135.53** | **Y304, Y309** |
| **Q9KTY1** | **Iron-sulfur cluster assembly scaffold protein IscU** | **1.04e+005** | **24.48** | **Y121** |
| **Q9KSW5** | **Isocitrate dehydrogenase [NADP]** | **2.68e+006** | **141.17** | **Y119** |
| **Q9KU60** | **Lysine--tRNA ligase** | **6.31e+005** | **86.81** | **Y112** |
| **Q9KPE6** | **Nicotinate-nucleotide pyrophosphorylase_ carboxylating** | **5.62e+004** | **17.37** | **Y13** |
| **Q9KT14** | **Oligopeptide ABC transporter_ periplasmic oligopeptide-binding protein/SBP_bac_5 domain-containing protein** | **1.26e+006** | **159.09** | **Y492** |
| **Q9KTX5** | **Peptidase B** | **5.71e+005** | **91.77** | **Y265** |
| **Q9KVH5** | **Peptide ABC transporter_ periplasmic peptide-binding protein/SBP_bac_5 domain-containing protein** | **4.29e+005** | **85.48** | **Y167** |
| **Q9KUA3** | **Peptide ABC transporter_ periplasmic peptide-binding protein** | **2.10e+006** | **201.28** | **Y48** |
| **Q9KR12** | **Peptidoglycan-associated protein** | **1.68e+005** | **24.68** | **Y116** |
| **Q9KSN6** | **Phenylalanine--tRNA ligase beta subunit** | **3.86e+005** | **126.85** | **Y744** |
| **Q9KM40** | **PhnA protein** | **1.38e+005** | **43.58** | **Y182** |
| **Q9KT08** | **Phosphate acetyltransferase** | **2.61e+006** | **222.58** | **Y663** |
| **Q9KNK0** | **Phosphoenolpyruvate carboxykinase (ATP)** | **5.21e+006** | **368.03** | **Y302, Y382** |
| **P0C6Q3** | **Phosphoglycerate kinase** | **5.40e+006** | **250.69** | **Y129** |
| **Q06951** | **Phosphomannomutase** | **3.51e+005** | **72.92** | **Y337** |
| **Q9KVS8** | **Phosphomethylpyrimidine synthase** | **8.03e+004** | **36.88** | **Y23** |
| **Q9KU76** | **Polyribonucleotide nucleotidyltransferase** | **8.21e+005** | **170.86** | **Y259** |
| **Q9KTM7** | **Proline--tRNA ligase** | **4.76e+005** | **117.86** | **Y133** |
| **Q9KNS8** | **Protein-export protein SecB** | **8.00e+005** | **94.27** | **Y18** |
| **Q9KPM0** | **Purine nucleoside phosphorylase DeoD-type 1** | **3.49e+005** | **62.20** | **Y161** |
| **Q9KNB2** | **Purine nucleoside phosphorylase DeoD-type 2** | **6.84e+005** | **97.68** | **Y161** |
| **Q9KS36** | **Putrescine-binding periplasmic protein** | **3.73e+005** | **65.45** | **Y60, Y99, Y107** |
| **Q9KPF4** | **Pyruvate dehydrogenase E1 component** | **3.07e+006** | **354.93** | **Y202, Y393,Y687, Y696, Y749, Y820** |
| **Q9KUN0** | **Pyruvate kinase** | **2.44e+006** | **260.43** | **Y40, Y110, Y390** |
| **Q9KLW8** | **Transaldolase** | **2.28e+006** | **185.35** | **Y217** |
| **Q9KUP2** | **Transketolase 1** | **3.54e+006** | **277.92** | **Y408** |
| **Q9KN05** | **Tryptophanase** | **2.45e+006** | **199.20** | **Y93, Y236** |
| **Q9KQ59** | **TyrA protein** | **3.49e+005** | **74.01** | **Y129, Y264** |
| **Q9KLK2** | **UPF0265 protein VC_A0741** | **2.68e+005** | **56.62** | **Y44** |
