## Supplementary Table 2 for "In-vivo protein nitration and de-nitration facilitate *Vibrio cholerae* cell survival under anaerobic condition: Consequences of Nitrite induced protein nitration"

**Table S2.** List of identified nitrated proteins of *Vibrio cholerae* strain C6706

| **Accession No.** | **Description** | **Average Normalized Abundances** | **Score** | **Nitration position** |
| --- | --- | --- | --- | --- |
| **A0A6M4VVP8** | **60 kDa chaperonin** | **4.58E+06** | **295.5** | **Y203** |
| **A0A6M4VVM5** | **Phosphoenolpyruvate carboxykinase (ATP)** | **3.47E+06** | **226.21** | **Y438** |
| **A0A6M4VTT4** | **Pyruvate dehydrogenase E1 component** | **3.59E+05** | **179.57** | **Y679, Y696** |
| **A0A6M4W100** | **Peptide ABC transporter substrate-binding protein** | **2.79E+06** | **174.63** | **Y139, Y142, Y250** |
| **A0A6M4VYC5** | **Aconitate hydratase B** | **1.13E+06** | **160.84** | **Y555, Y556** |
| **A0A6M4VYZ9** | **Fructose-bisphosphate aldolase** | **3.35E+05** | **120.2** | **Y302, Y309** |
| **A0A6M4VUL7** | **Di-hydro lipoyl lysine-residue succinyl transferase component of 2-oxoglutarate dehydrogenase complex** | **9.12E+05** | **118.04** | **Y219, Y248** |
| **A0A6M4WBE2** | **Tryptophanase** | **2.31E+06** | **107.57** | **Y109, Y456** |
| **A0A6M4VTK5** | **Glutamine synthetase** | **3.78E+05** | **97.19** | **Y398, Y421** |
| **A0A6M4W5L2** | **Glycerol kinase** | **5.01E+04** | **96.77** | **Y99** |
| **A0A6M4VVL9** | **Catalase-peroxidase** | **1.37E+05** | **89.71** | **Y329, Y331, Y336** |
| **A0A6M4VU98** | **OmpA family protein** | **1.07E+05** | **87.16** | **Y237** |
| **A0A6M4W154** | **Transketolase** | **1.11E+05** | **86.2** | **Y408** |
| **A0A6M4VZG8** | **ATP synthase subunit alpha** | **4.43E+05** | **76.27** | **Y192, Y195** |
| **A0A6M4VXE1** | **Porin OmpT** | **2.62E+05** | **70.7** | **Y33, Y171, Y272, Y289** |
| **A0A6M4W4Q0** | **Amino acid ABC transporter substrate-binding protein** | **4691.99** | **57.9** | **Y37** |
| **A0A6M4VZ77** | **GDP-mannose 4_6-dehydratase** | **6.77E+04** | **57.1** | **Y258, Y312** |
| **A0A6M4W1X8** | **Oxoglutarate dehydrogenase (succinyl-transferring)** | **5.91E+04** | **52.06** | **Y567** |
| **A0A6M4W0H5** | **MipA/OmpV family protein** | **2.29E+04** | **50.07** | **Y214** |
| **A0A6M4VYD3** | **Lipoprotein** | **5.12E+04** | **49.61** | **Y225** |
| **A0A6M4W2K7** | **Peptidoglycan-associated protein** | **2095.67** | **48.74** | **Y131** |
| **A0A6M4VUK3** | **50S ribosomal protein L5** | **1.37E+05** | **44.38** | **Y7, Y8** |
| **A0A6M4W6S7** | **50S ribosomal protein L11** | **1.95E+06** | **41.1** | **Y8** |
| **A0A6M4VYH8** | **Universal stress protein UspE** | **3264.57** | **38.86** | **Y81** |
| **A0A6M4W3M6** | **Uncharacterized protein** | **1473.15** | **37.13** | **Y189, 198** |
| **A0A6M4VZ01** | **Rsd/AlgQ family anti-sigma factor** | **1.47E+04** | **36.22** | **Y61** |
| **A0A6M4VY61** | **Adenylate kinase** | **9.32E+04** | **35.86** | **Y133, Y181, Y182** |
| **A0A6M4VVE9** | **Universal stress protein** | **9.31E+04** | **33.82** | **Y79** |
| **A0A6M4W0X6** | **Stringent starvation protein A** | **2722.62** | **26.22** | **Y92** |
| **A0A6M4VWW9** | **Succinate--CoA ligase [ADP-forming] subunit alpha** | **4.78E+04** | **21.14** | **Y31** |
