## Supplementary Table 3 for "In-vivo protein nitration and de-nitration facilitate *Vibrio cholerae* cell survival under anaerobic condition: Consequences of Nitrite induced protein nitration"

**Table S3.** Pathway wise distribution of various nitrated proteins identified N16961 and C6706 strain of *Vibrio cholera*e

| **Type** | **N16961** | **C6706** |
| --- | --- | --- |
| **Glycolysis/Gluconeogenesis**  **Or PPP** | - **2_3-bisphosphoglycerate-independent phosphoglycerate mutase** - **Acetyltransferase component of pyruvate dehydrogenase complex** - **Dihydrolipoyl dehydrogenase** - **Enolase** - **Glyceraldehyde-3-phosphate dehydrogenase** - **Phosphoenolpyruvate carboxykinase (ATP)** - **Phosphoglycerate kinase** - **Pyruvate dehydrogenase E1 component** - **Pyruvate kinase** - **Transaldolase** - **Transketolase 1** | - **Phosphoenolpyruvate carboxykinase (ATP)** - **Pyruvate dehydrogenase E1 component** - **Fructose-bisphosphate aldolase** - **Transketolase** |
| **TCA cycle** | - **Citrate synthase** - **Formate acetyltransferase** - **Isocitrate dehydrogenase [NADP]** | - **Aconitate hydratase B** - **Oxoglutarate dehydrogenase (succinyl-transferring)** - **Succinate--CoA ligase [ADP-forming] subunit alpha** |
| **Electron Transport Chain** | - **Succinate dehydrogenase flavoprotein subunit** - **ATP synthase subunit beta** - **ATP synthase subunit alpha** | - **ATP synthase subunit alpha** |
| **Nucleotide metabolism** | - **2`_3`-cyclic-nucleotide 2`-phosphodiesterase** - **CTP synthase** - **Phosphomethylpyrimidine synthase** - **Polyribonucleotide nucleotidyltransferase** - **Purine nucleoside phosphorylase DeoD** |  |
| **Carbohydrate metabolism** | - **Alpha-1_4 glucan phosphorylase** - **GDP-mannose 4_6-dehydratase** - **Glucose-1-phosphate adenylyltransferase 1** - **Phosphomannomutase** | - **GDP-mannose 4_6-dehydratase** |
| **Amino acid/Protein metabolism** | - **Peptidyl-prolyl cis-trans isomerase** - **Asparagine--tRNA ligase** - **Aspartate ammonia-lyase** - **Glutamine synthetase** - **Lysine--tRNA ligase** - **Peptidase B** - **Phenylalanine--tRNA ligase beta subunit** - **Proline--tRNA ligase** - **Tryptophanase** - **TyrA protein** | - **Di-hydro lipoyl lysine-residue succinyl transferase component of 2-oxoglutarate dehydrogenase complex** - **Tryptophanase** - **Glutamine synthetase** |
| **Lipid metabolism** | - **2-dehydro-3-deoxyphosphooctonate aldolase** - **Acetyl-coA carboxylase carboxyl transferase subunit alpha** - **Glycerol kinase** | - **Glycerol kinase** - **Lipoprotein** |
| **Porin or transporter proteins** | - **Oligopeptide ABC transporter_ periplasmic oligopeptide-binding protein/SBP_bac_5 domain-containing protein** - **Peptide ABC transporter_ periplasmic peptide-binding protein** - **Protein-export protein SecB** - **Putrescine-binding periplasmic protein** | - **Peptide ABC transporter substrate-binding protein** - **OmpA family protein** - **Porin OmpT** - **Amino acid ABC transporter substrate-binding protein** - **MipA/OmpV family protein** |
| **Redox/stress protective proteins** | - **Thioredoxin Reductase** - **Aldehyde-alcohol dehydrogenase** - **Glutathione Reductase** - **Iron-sulfur cluster assembly scaffold protein IscU** | - **Catalase-peroxidase** - **Universal stress protein UspE** - **Stringent starvation protein-A** |
| **Other housekeeping proteins/enzymes** | - **50S ribosomal protein L21** - **Adenylate kinase** - **DNA-directed RNA polymerase subunit beta (rpoB)** - **HTH-type transcriptional repressor PurR** - **Iron-containing alcohol dehydrogenase family protein RfbM** - **Nicotinate-nucleotide pyrophosphorylase_ carboxylating** - **Peptidoglycan-associated protein** - **PhnA protein** - **Phosphate acetyltransferase** - **60 kDa chaperonin** - **Elongation factor G 2** | - **60 kDa chaperonin** - **Peptidoglycan-associated protein** - **50S ribosomal protein L5** - **50S ribosomal protein L11** - **Rsd/AlgQ family anti-sigma factor** - **Adenylate kinase** |
